# Comparing Phylogeographies: Incompatible Geographical Histories in Pathogens’ Genomes

**DOI:** 10.1101/2020.01.10.902460

**Authors:** Benjamin Singer, Antonello Di Nardo, Jotun Hein, Luca Ferretti

**Author notes:** These authors contributed equally (Benjamin Singer, Antonello Di Nardo).

## Abstract

Modern phylogeography aims at reconstructing the geographic diffusion of organisms based on their genomic sequences and spatial information. Phylogeographic approaches usually ignore the possibility of recombination, which decouples the evolutionary and geographic histories of different parts of the genome. Genomic regions of recombining or reassorting pathogens often originate and evolve at different times and locations, which characterised their unique spatial histories. Measuring the extent of these differences requires new methods to compare geographic information on phylogenetic trees reconstructed from different parts of the genome. Here we develop for the first time a set of measures of *phylogeographic incompatibility* aimed at detecting differences between geographical histories in terms of distances between phylogeographies. We study the effect of varying demography and recombination on phylogeographic incompatibilities using coalescent simulations. We further apply these measures to the evolutionary history of human and livestock pathogens, either reassorting or recombining, such as the Victoria and Yamagata lineages of influenza B and the O/Ind-2001 foot-and-mouth disease virus strain. Our results reveal diverse geographical paths of diffusion that characterise the origins and evolutionary histories of different viral genes and genomic segments. phylogeography, recombination, viral evolution

## 1 Introduction

The study of the evolutionary biology and phylodynamics of pathogens has made significant contributions to disease control over the last few decades [Grubaugh et al., 2019, Baele et al., 2018]. Pathogens are dispersed with their human and animal hosts, both of which are seeing increasing mobility as an effect of globalisation and the access of emerging economies to international markets. This condition offers an increasing opportunity for some pathogens to spread rapidly at the global level, influenza being one of the best known examples [Lemey et al., 2014]. The elevated mobility pattern combines with high evolution rates in some pathogens to couple the processes of evolution and spatial diffusion in these organisms. To understand the emergence, adaptation and spread of pathogens, therefore, it is possible to use the methods provided by the field of modern phylogeography, that jointly infer the evolutionary history (i.e. phylogeny) and the spatial history (i.e. geographic diffusion) of an organism [Dudas et al., 2017].

Phylogeographic methods can make use of the mathematical representation of a phylogeny as a tree, allowing the spatial evolution from the root to the tips to be seen as a diffusion or migration process. This diffusion process proceeds independently on each branch of the tree, with a single origin at the root. The resulting structure greatly simplifies mathematical and computational treatments of systems composed of coupled evolution and migration processes. However, the phylogenetic tree of a set of DNA or protein sequences contains information not only on the relatedness of sequences, but also on their evolutionary history. There exist many different methods for reconstructing such a tree. If genome isolates have been collected from different places, their history could be affected by the geographic structure of the populations, including the rates of migration across different locations [Lemey et al., 2014]. The first step to understanding these effects is the reconstruction of a phylogenetic tree with geographical information attached to the internal nodes, obtained through reconstruction of the spatial movements of lineages in the tree. Such an object corresponds to the phylogeographic history of the given set of sequences. For simplicity, in this paper we will refer to it as a *phylogeography*.

A range of techniques have been developed for the reconstruction of phylogeographies [Faria et al., 2011]. These can be classified according to how the migration process of ancestral lineages is modelled on the tree: either in the form of instantaneous diffusion of discrete traits (such as finite spatial locations), i.e. discrete phylogeography [De Maio et al., 2015, Kühnert et al., 2016, Lemey et al., 2009, Müller et al., 2017]; or as relaxed random walk diffusion in two-dimensional geographic space, i.e. continuous phylogeography [Bouckaert, 2016, Dellicour et al., 2016, Gill et al., 2017, Lemey et al., 2010]. Many of these approaches focus on a single tree, obtained from concatenation based phylogeny or selected sections of a genome, and therefore do not take into account the decoupling of the evolutionary history of different genomic regions due to recombination and/or related processes.

For many species of eukaryotes, as well as for some viruses and bacteria, phylogenies are not tree-like. In the presence of recombination, reassortment or horizontal gene transfer, different regions of the genome might have potentially evolved from unique ancestors. This decouples the evolutionary histories of different genes or regions of the genome, resulting in trees with cross-branch reticulations if the number of recombination events is small, or in phylogenetic networks if recombination is widespread. Recombination leads to organisms that inherit genomic fragments from multiple parent lineages, causing phylogenetic inference methods applied to different parts of the genome to give rise to incompatible trees. By applying a measure of tree distance to these trees, it is possible to make certain claims about the amount of recombination in the ancestral population [Chung et al., 2013], for instance by setting a lower bound on the number of recombination events between loci. Any non-zero tree distance reveals an incompatibility. If the genealogies of different parts of the same genome are incompatible, these incompatibilities can be evidence of recombination, horizontal gene transfer, reassortment, or gene conversion. A variety of incompatibility measures and tree distances exist for phylogenies, most notably the well-known Robinson-Foulds metric and the family known as tree edit (or Levenshtein) distances [Day, 1984], including the subtree prune and regraft (SPR) distance [Semple and Steel, 2003].

Recombination can lead to a decoupling of geographic histories between loci, just as it can lead to a decoupling of phylogentic histories. The resulting ensemble of spatial trajectories may reveal functional and evolutionary features of different genes. However, phylogeographic inference is highly non-trivial in the presence of reticulations. Reticulations form phylogenetic loops, meaning that two lineages can split at some time in the past, move in geographic space, then converge to the same location and combine into a single lineage. They can share pieces of genetic material with different geographical origins, or copies of material which have reached their current geographical distribution and evolutionary trajectory in different ways. This means that the spatial evolution of a given pair of lineages can no longer be assumed independent, and some of the advantages of working with diffusion processes on a tree are lost.

The purpose of this paper is to propose and define two mathematical measures of the degree of difference between reconstructed phylogeographies arising from different loci. Only one of these is a true distance in the sense of providing a metric on a vector space, so we will use the more general term ‘incompatibility measure’. A non-zero value of an incompatibility measure between two phylogeographies implies that they are incompatible, and is evidence of both recombination and different patterns of spatial diffusion between loci. Comparing incompatibilities between the phylogeographies of different loci can reveal which regions tend either not to recombine much, or to remain associated during spatial diffusion and those which, conversely, have highly divergent spatial histories. We apply these measures of phylogeographic incompatibility to reconstructed histories of a reassorting human pathogen, the influenza B virus, as well as a recombining animal pathogen, the foot-and-mouth disease virus (FMDV).

## 2 New Approaches

Any measure of phylogeographic incompatibility will inevitably be related to phylogenetic distances, due to the mathematical relation between phylogeographies and phylogenies. In this section we outline our approach to calculating phylogeographic incompatibilities, starting with a comparison to the phylogenetic case and detailing two possible ways to incorporate geographic information. We then provide an approach that accounts for uncertainties in phylogeographic reconstruction.

### 2.1 Phylogenetic incompatibilities

Many different measures of tree distance/incompatibility have been developed in the past. In most comparisons, we will use only the two measures most closely related to the phylogeographic incompatibilities introduced in the next sections.

The first is the path distance measure (hereinafter referred to as “path”) introduced by [Steel and Penny, 1993]. This measure compares the heights of the most recent common ancestors (MRCAs) of each pair of leaves for the two trees and computes the norm of their differences. Since most pairwise MRCAs are localised close to the root, the path distance is strongly affected by topological variation of the upper part of the tree.

The second is the natural distance based on the size of the maximum agreement sub-tree (“MAST”) between the two trees [Steel and Warnow, 1993]. The MAST estimate is very large for similar trees, while it is computed using fewer and fewer leaves as the dissimilarity between the trees increases. The MAST distance is dramatically sensitive to changes near the root but it is also generally sensitive to variability across the whole tree. Note that the normalised MAST distance is usually defined as the fraction of tips that do *not* belong to the MAST.

We also consider four other distance measures defined on phylogenies (without a geographical element):

- Robinson-Fould metric (“RF”) [Robinson and Foulds, 1981]
- Robinson-Fould metric weighted by branch lengths (“WRF”) [Robinson and Foulds, 1981]
- subtree-prune-and-regraft edit distance (“SPR”) [De Oliveira Martins et al., 2014, Allen and Steel, 2001]
- Kuhner-Felsenstein branch score distance (“KF”) [Kuhner and Felsenstein, 1994]

Each of these incompatibility measures defines a proper metric in the space of rooted binary trees with branches of finite length. However, all these distances capture different ways in which phylogenies can differ. Given the high-dimensional and heterogeneous nature of phylogenetic spaces, it is not surprising that there is little agreement among the patterns of distances, as illustrated in the Appendix. As an example, trees differing by the topology of the upper half (near the root) tend to have a high MAST distance and an comparatively even larger path distance; the opposite is true for trees differing in the topology of the lower half (near the tips), which tend to be close in terms of path distance but differ sensibly in terms of MAST distance.

Throughout this paper, we used the version of these measures adapted to rooted binary trees. These measures are currently implemented in the R packages *APE* [Paradis and Schliep, 2018] and *phangorn* [Schliep, 2010].

### 2.2 Phylogeographic space

The first step in building natural methods of comparison for phylogeographies is to define what a phylogeography is and to situate it in a well-defined mathematical space. The incompatibility measures should facilitate the definition of distance measures over this space. Such a space needs to be able to represent both phylogenetic and geographical information.

We define a phylogeography as a tree with geographical discrete information assigned to each node of the tree. For practical purposes, inferred phylogenies are represented by rooted phylogenetic trees with branch lengths and labelled tips. Geographical space can be represented either as a network of discrete locations or as a continuous surface. Our approach covers both cases. The question then becomes how to combine these spaces, and how to define a joint measure on the resulting combination. These two questions are not independent, as different spaces require different approaches for comparing phylogeographies.

We present a simple definition of phylogeographic space. The most general space is denoted by _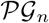_, with integer *n*, and its elements are rooted binary trees with *n* labelled leaves and with locations assigned to all nodes (both leaves and internal nodes). The illustrations in figure 1 are of phylogeographies of this kind. This space corresponds to the Cartesian product of the space of bifurcating rooted trees with *n* leaves 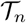 and the Cartesian power of the geographic space 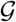 to the number of nodes, i.e. 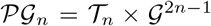. In practice, we will work on subspaces of 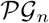 with a given vector of tip locations **g**, since the locations of the leaves are usually known and given as input into phylogeographic analyses. For a given set of locations **g**, the corresponding phylogeographic subspace 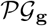 is isomorphic to 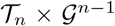. In the following, when discussing 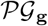 for a specific (although implicit) choice of **g**, we will refer to it simply as 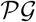.

**Figure 1:**
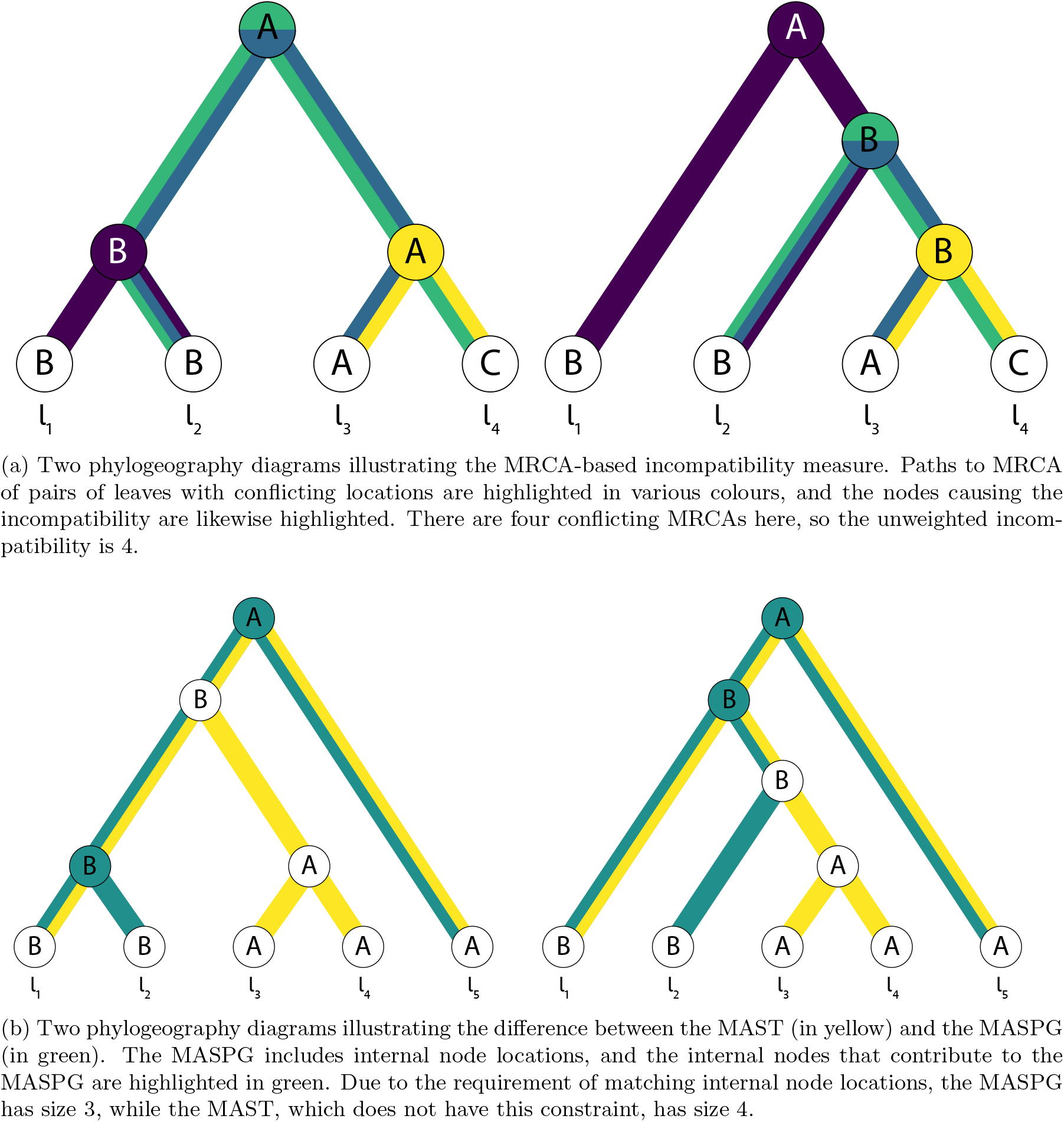
An illustrations of the two measures of phylogeographic incompatibility presented in this paper:(a) pairwise MRCA incompatibility, (b) Maximum Agreement PhyloGeography (MASPG).

### 2.3 Pairwise MRCA Incompatibilities

The geographic space 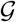 has a metric 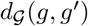 already defined on it. We can use this distance to define a purely geographic measure of incompatibilities on 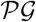. However, this requires a pairwise comparison of spatial locations. Since locations are assigned to nodes on different trees, we must solve the matching of nodes across trees with different shapes. The simplest approach is to map the MRCAs for pairs of leaves, i.e. compare nodes which are the MRCA of the same pair of leaves on different trees.

The first step to performing such a comparison for a pair of trees with *n* leaves is to project thephylogeography *PG* to a 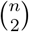-ary vector **g**_*MRCA*_(*PG*), where the element *g*_*MRCA*_(*l*_1_, *l*_2_) is the location of the MRCA of the tips *l*_1_ and *l*_2_. Such vectors span a subspace of the 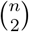-ary Cartesian power of 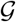. The *L*_1_ product metric is a natural metric for this space and therefore it can be used to compute geographic incompatibilities between phylogeographies.

The incompatibility measure between two phylogeographies *PG*_1_ and *PG*_2_ is then defined as

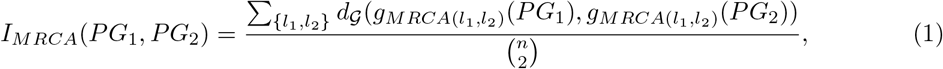

where {*l*_1_, *l*_2_} range over the set of pairs of leaves, *g*_*MRCA*_(*l*_1_, *l*_2_)(*PG*_*k*_) is the location assigned to the MRCA of leaves *l*_1_ and *l*_2_ in the phylogeography *PG*_*k*_, and 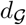 is the metric on the geographic network. The normalisation factor 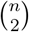 ensures that the minimum and maximum possible values of the measure are 0 and 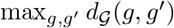, respectively.

For a simple example of how to calculate *I*_*MRCA*_, consider the phylogeographies in Figure 1a. Take *A*, *B*, and *C* to refer to the arbitrary locations Argentina, Bolivia and Chile, and place these countries on a network where there is a pairwise distance of 1 between each pair, since they all share borders. The MRCA vectors are therefore *g*_*MRCA*_(*PG*_1_) = (*B, A, A, A, A, A*) for the phylogeography on the left, and *g*_*MRCA*_(*PG*_2_) = (*A, A, A, B, B, B*) for the phylogeography on the right. So the total incompatibility is 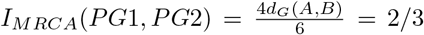. Here, the incompatibility comes about due to four comparisons between Argentina and Bolivia, and the only MRCA paths that do not contribute to the incompatibility are between *l*_1_ and *l*_3_, and between *l*_1_ and *l*_4_, which meet at the root in both phylogeographies.

*I*_*MRCA*_ is especially sensitive to the locations assigned to nodes with many descendants, the strongest example of this being the root of the tree, which can contribute to half or more of the comparisons made in calculating the incompatibility. This sensitivity depends in part on the tree shape, with the root contributing more in symmetric trees than in asymmetric ones. This is because leaf pairs are more likely to find their MRCA at the root when the tree is more symmetric, and more likely to find their MRCA on the trunk of the tree when the tree is less symmetric.

Note that *I*_*MRCA*_ is not a distance on 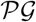, since trees with a different topology may still have no incompatibilities. However, this issue is easily solved. In fact, any positive linear combination of *I*_*MRCA*_ and a tree distance on 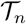 is a proper metric on 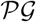.

### 2.4 Maximum Agreement Sub-Phylogeography

To incorporate both phylogenetic and geographic differences into a metric i n a unified way, an alternative approach is to extend an existing metric on 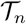 into 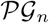. One of the simplest measures of phylogenetic incompatibility is based on the notion of a maximum agreement subtree, which we will here extend to phylogeographies.

As other phylogenetic distances, this accounts for the similarity between different trees based on shared subtrees. There already exists a fairly comprehensive body of literature on finding MASTs between pairs of trees [Hein et al., 1996, Steel, 2016, Semple and Steel, 2003]. An agreement subtree between the trees *T*_1_ and *T*_2_ with labelled leaves is a tree *S*, with labelled leaves, such that the subtrees induced on *T*_1_ and *T*_2_ by the set of leaves corresponding to those in *S* are both identical to *S*. The MAST is the largest such subtree. Figure 1b provides an example of a MAST.

We can use the MAST to define a metric over 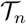. To find the distance between *T*_1_ and *T*_2_, we start to find *MAST*(*T*_1_, *T*_2_), and then write *d*_*MAST*_ (*T*_1_, *T*_2_) = *n* − |*MAST*(*T*_1_, *T*_2_)|. Here, |*T*| is the number of leaves on *T*, and *n* = |*T*_1_| = |*T*_2_|. Thus, for the trees in the example in Figure 1b, *d*_*MAST*_(*T*_1_, *T*_2_) = 1, since the trees are of size five and the MAST i s of size four.

Now we extend this distance to a metric on 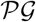. First, we define a maximum agreement sub-phylogeography (MASPG). Take two phylogeographies *PG*_1_ and *PG*_2_, with locations assigned to each node. Let us define an agreement sub-phylogeography as a sub-phylogeography without unary nodes such that (i) the corresponding subtree is an agreement subtree and (ii) for each node in the subtree, the location assigned to the node matches the locations on the corresponding subtrees in both *PG*_1_ and *PG*_2_. That is, the induced subtrees must be identical not only in topology but also in the geographical assignment of the nodes. The MASPG is defined as the largest such sub-phylogeography.

We can then define the phylogeographic incompatibility between two phylogeographies *PG*_1_ and *PG*_2_, both with leaf number *n* = |*PG*_1_| = |*PG*_2_|, as

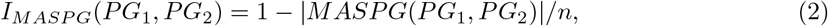

where |*T*| is the number of leaves on the tree *T*. With this normalisation, the minimum and maximum possible values of *I*_*MASPG*_ are 0 and 1, respectively. For the phylogeographies in figure 1b the MASPG is of size three and the phylogeographies are of size five, so the incompatibility is 0.4. *I*_*MASPG*_ is a metric on 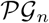, as shown in Appendix 1.

### 2.5 Incompatibility between distributions of phylogeographies

In practical applications, phylogeographies are not known with absolute precision, but they are inferred from noisy and biased data (e.g. biased sampling of subpopulations). In fact, Bayesian approaches to phylogeographic inference actually return a posterior distribution of phylogeographies. For this reason, we consider the more general case of comparisons between two *distributions* of phylogeographies. In such a case, the objects of comparison are two random variables PG_1_ and PG_2_ derived from different loci along a genome. These variables are likely to be highly correlated even if their posterior (marginal) distributions are inferred independently, due to linkage between loci and the fixed assignment of tip locations from sample metadata. This causes strong biases if the expected value of *I*_*MRCA*_ is computed assuming the two distributions as independent.

The simplest and most conservative choice is to compute the incompatibility as a Wasserstein metric (or earth mover’s distance) between the distributions, using one of the incompatibility measures proposed above as cost function. This choice reduces the likelihood of spurious incompatibilities due to correlations between the inferred phylogeographies at different loci.

Hence, we can use *I*_*MRCA*_ or *I*_*MASPG*_ as a cost function for differences between phylogeographies, yielding a definition of incompatibility between PG_1_ and PG_2_ as the first Wasserstein metric on the corresponding distributions. In practice, we estimate this by taking *N* samples from each posterior, pairing these samples in all possible *N* ! ways (e.g. *s*_1_ with 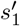, *s*_2_ with 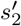 etc), computing the mean of the incompatibilities 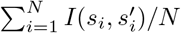 between all pairs of samples from the two distributions, then minimising this quantity over all choices of pairings.

## 3 Results

### 3.1 Effect of evolutionary parameters on phylogeographic incompatibilities

To understand the impact of recombination and migration rates on phylogeographic incompatibilities, we generated random phylogeographies from structured coalescent models with three populations/locations. We simulated both coalescent trees and node locations of 20 sequences from each location, all sampled at the present time. Simulations were performed by varying population-scaled recombination rate *ρ* and migration rate *μ*, as well as migration patterns.

Incompatibilities are generated by different mechanisms for the patterns of migration represented e.g. in the models in Figure 2. In model (iii), there is only one possible chain of geographical transmissions for each lineage, but incompatibilities are generated by the different pace of lineages along this chain. These differences originate from neutral (stochastic) or selective differences in the patterns of coalescence or migration of the lineages. In model (iv), multiple patterns of transmission represent another important contribution to phylogeographic incompatibilities, with lineages following different routes between the source and the final location. In model (i), an additional source of incompatibility is provided by bidirectional spread, with lineages bouncing back and forth or wandering between distant locations.

**Figure 2:**
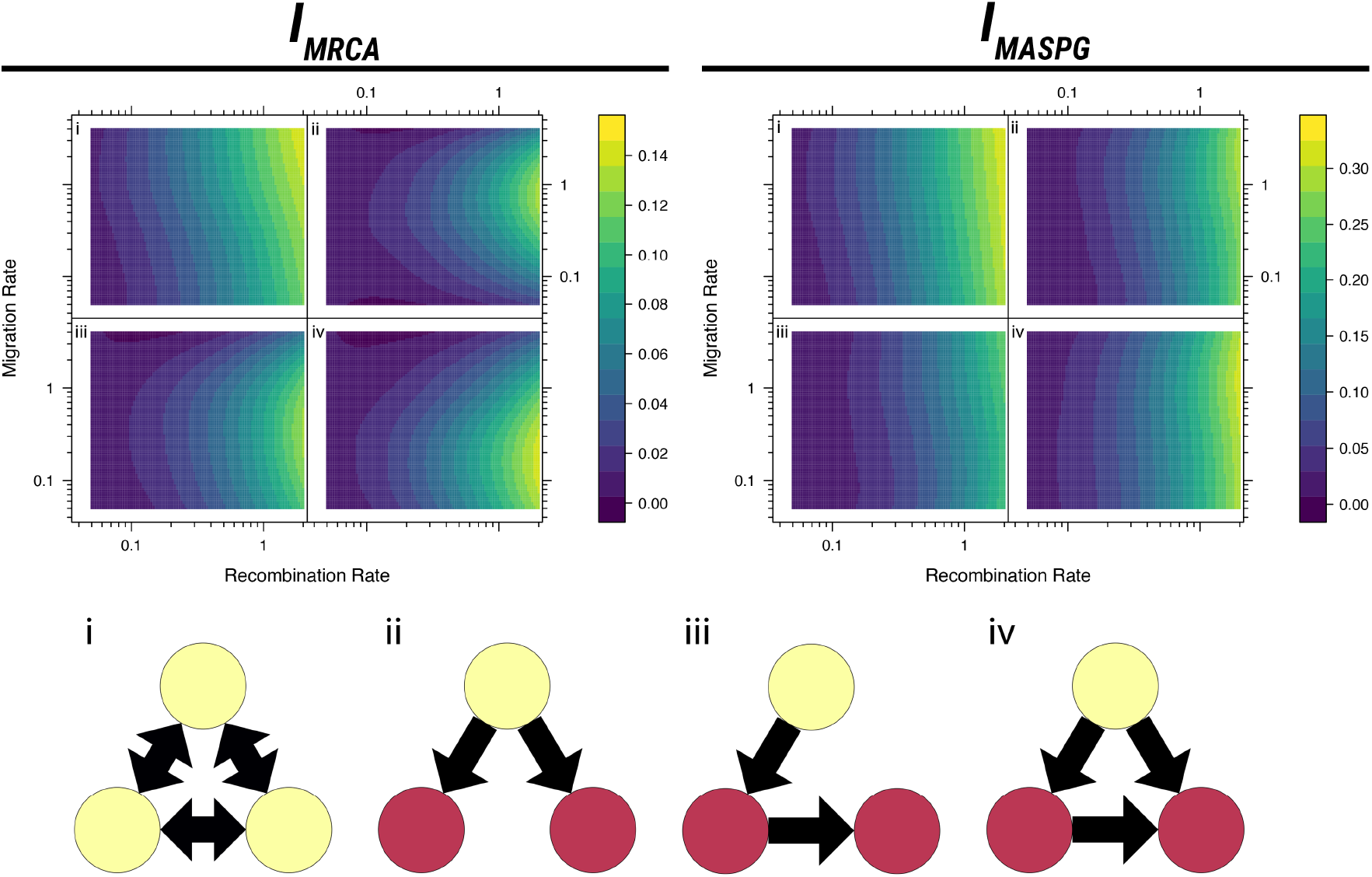
Effect of evolutionary parameters on phylogeographic incompatibilities, using 100 simulations at each calculated pair of parameter values. The plots show smoothed surfaces of each phylogeographic incompatibility measure, labelled with the corresponding network structure. The network structures are illustrated with locations in which the population can originate coloured in yellow.

Figure 2 reveals that the incompatibility between simulated trees increases with both recombination rate and migration rate, at least for low/moderate rates of migration. This is to be expected, since increased recombination implies more different trees, while low migration leads to less spatial transitions along branches and therefore more similar geographic histories.

In most models, changes in recombination rates have a much stronger impact on phylogeographic incompatibilities than changes in migration rates. This can be explained by the fact that recombination is necessary to actually observe a difference in migration histories. Without any topological variation of the tree, the phylogeographic reconstruction would not differ and any change in migration patterns would be undetectable. In fact, incompatibilities are very weak for recombination rates *ρ* < 1 and increase steeply for larger values of *ρ*. Incompatibilities tend to be weaker as well for low migration rates *μ* ≪ 1.

As outlined above, migration patterns have a major impact on incompatibilities, depending on the migration rate and the specific measure of incompatibility. Incompatibilities tend to be small if there is a single source population (Figure 2-ii), since this imposes strong constraints on the migration histories and on the origin of the ancestors of the sample. Scenarios with longer transmission chains (Figure 2-iii) or more complex patterns of transmission (Figure 2-iv) lead to a slight increase in differences among phylogeographies and a more complex dependence on migration rates. Finally, in scenarios with multiple bidirectional migration patterns, migration histories can be wildly different, hence incompatibilities tend to be much larger (Figure 2-i) and to increase monotonically but weakly with migration rates.

In some scenarios, different measures of incompatibilities show strikingly different trends. This is due to the fact that they capture different components of the signal. As an example, widespread differences in recent geographical history would give rise to a much stronger signal on MASPG than on MRCA incompatibilities.

### 3.2 Phylogeography of the O/ME-SA/Ind-2001 FMDV lineage epidemics

Foot-and-mouth disease is a viral disease affecting wild and domesticated cloven-hoofed animals [Alexandersen et al., 2003] and it is one of the most economically relevant diseases of livestock worldwide [Knight-Jones and Rushton, 2013]. It is caused by the FMDV, a positive-sense single-stranded RNA virus belonging to the family *Picornaviridae*. This virus has a relatively short (of about 8kb) but highly variable linear genome coding for 12 proteins with different functions [Jackson et al., 2003]. It provides an interesting case study for phylogeographic incompatibilities, since it has a non-trivial geographic structure, being spread out across host populations world-wide [Brito et al., 2017, Di Nardo et al., 2011], and a high recombination rate, especially between structural and non-structural proteins [Carrillo et al., 2005, Jackson et al., 2007].

Whole-genome sequences from a recently analysed FMDV lineage (the O/ME-SA/Ind-2001) were provided by the World Reference Laboratory for FMD (WRLFMD) at the Pirbright Institute [Bachanek-Bankowska et al., 2018]. Phylogeographic analyses were performed on most loci along the genome, as well as on the WGS. The phylogenetic and phylogeographic incompatibilities are illustrated in Figure 3.

**Figure 3:**
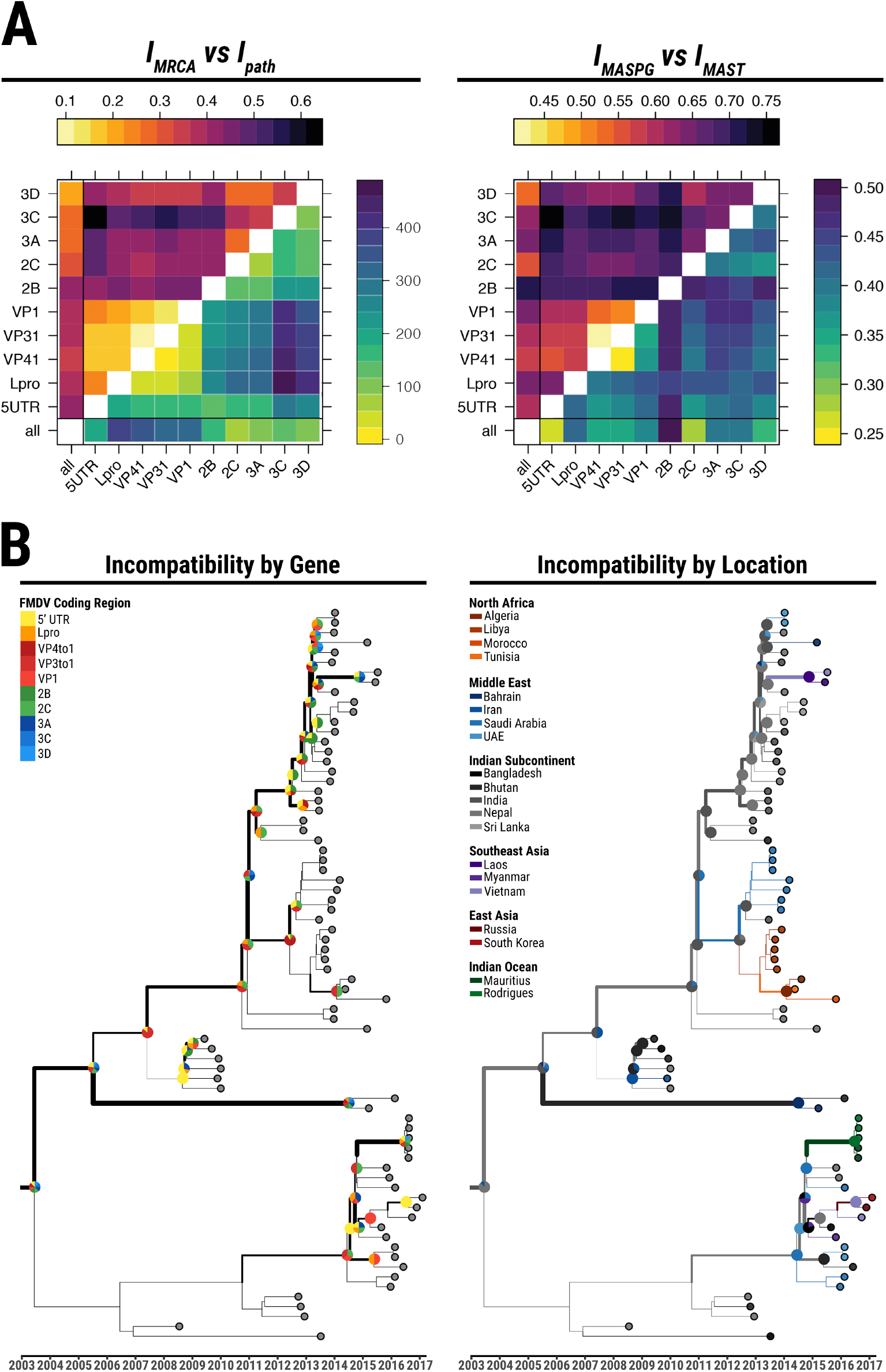
Phylogenetic and phylogeographic incompatibilities estimated for the FMDV O/ME-SA/Ind-2001 epidemics. A panel - Comparisons of various incompatibility measures between phylogeographies reconstructed from segments of the FMDV O/ME-SA/Ind-2001 genome. In the plot on the left, *I_MRCA_* (upper triangle) is compared to *I_path_* (lower triangle). In the plot on the right, *I_MASPG_* (upper triangle) is compared to IMAST (lower triangle). B panel - Illustration of the incompatibilities along the MCC tree. The width of each branch represents the total amount of phylogeographic MRCA incompatibilities between all trees built from individual genomic regions and the whole-genome tree. Pie charts represent the distribution of MRCA incompatibilities per genomic region (left) and per location (right) on the corresponding nodes. Branch colors on the right represent the phylogeography reconstructed from the whole-genome sequence alignment.

The two measures of phylogeographic incompatibility are only partially related. However, this is true for usual phylogenetic incompatibilities as well, as can be seen comparing the left and right panel of Figure 3.

Our results show that the portions of the genome coding for the capsid structure (comprising of the structural proteins from VP1 to VP4) have highly coupled histories, both phylogenetically and geographically. This fits with the well-known fact that this region exhibits a relatively low recombination rate [Carrillo et al., 2005, Jackson et al., 2007]. The non-structural genes at the 3’ end of the genome (i.e. 2C, 3A, 3C, and 3D) also share similar phylogenetic and geographic histories. The majority of incompatibilities are located along the tree trunk of the main sublineage, as illustrated in Figure 3: they are mostly related to differences in reconstructed virus movements within the Indian subcontinent, as well as variability in the timing of transmission of different genomic regions to the Middle-East.

Some genomic regions share similar phylogenies, but clearly distinct phylogeographies. This is the case e.g. for the 5’ UTR versus 2B-3A and for 2B versus 2C-3A, according to the MRCA measures estimated. In fact, topological similarity does not always reflect compatible geographical histories, and this relation is not reflected in the phylogenetic distances: for example, the WGS reconstruction is phylogenetically similar to the one from the L-protease, but the geographical histories are quite different.

It is important to note that the phylogeography created using the WGS does not provide a good summary of the phylogeographies of the individual genomic regions, and in fact it only resembles the phylogeographies from the genome segments comprising of the 2C and 3A proteins, which tend to be quite dissimilar to those of the rest of the sequences. This could be due to the effect produced on the WGS topology by recombinant viruses (i.e. the O/BAR/15/2015 and O/BHU/3/2016) [Bachanek-Bankowska et al., 2018], which provide the strongest contribution to incompatibilities (as is evident from the widest branch in Figure 3).

It is therefore important in FMDV evolutionary studies to take account of geographic as well as phylogenetic differences when comparing reconstructed evolutionary histories of distinct genomic regions outside of the capsid.

### 3.3 Spatial evolution of genomic segments of Influenza B

Influenza B virus is a segmented virus in the family *Orthomyxoviridae*. It is a major human pathogen, although it causes a lesser threat to public health than the related influenza A virus. Influenza B virus has a short positive-sense single-stranded RNA genome (of about 15 kb) organised in eight linear segments [Bouvier and Palese, 2008]. Reassortment of these segments plays a major role in the evolution of the virus, hence influenza B represents an interesting case study for phylogeographic incompatibilities in reassorting pathogens. Two main lineages (denoted as Victoria and Yamagata) are recognised based on differences in the hemagglutinin protein. It has been previously shown how these lineages evolved under different dynamics [Langat et al., 2017].

We analysed a set of 242 sequences (122 of the Victoria lineage, and 120 of the Yamagata) from a recent study of worldwide evolution of these lineages [Bedford et al., 2015]. These sequences include 5 genes (HA, NA, PB1 and PB2, and NS1 protein) located on different segments and therefore reassorting freely among them. Phylogenetic and phylogeographic incompatibilities among these segments are illustrated in Figure 4 for both lineages. We also reconstructed and compared the phylogeographic history of the joint sequences of all these genes (8.6 kb).

**Figure 4:**
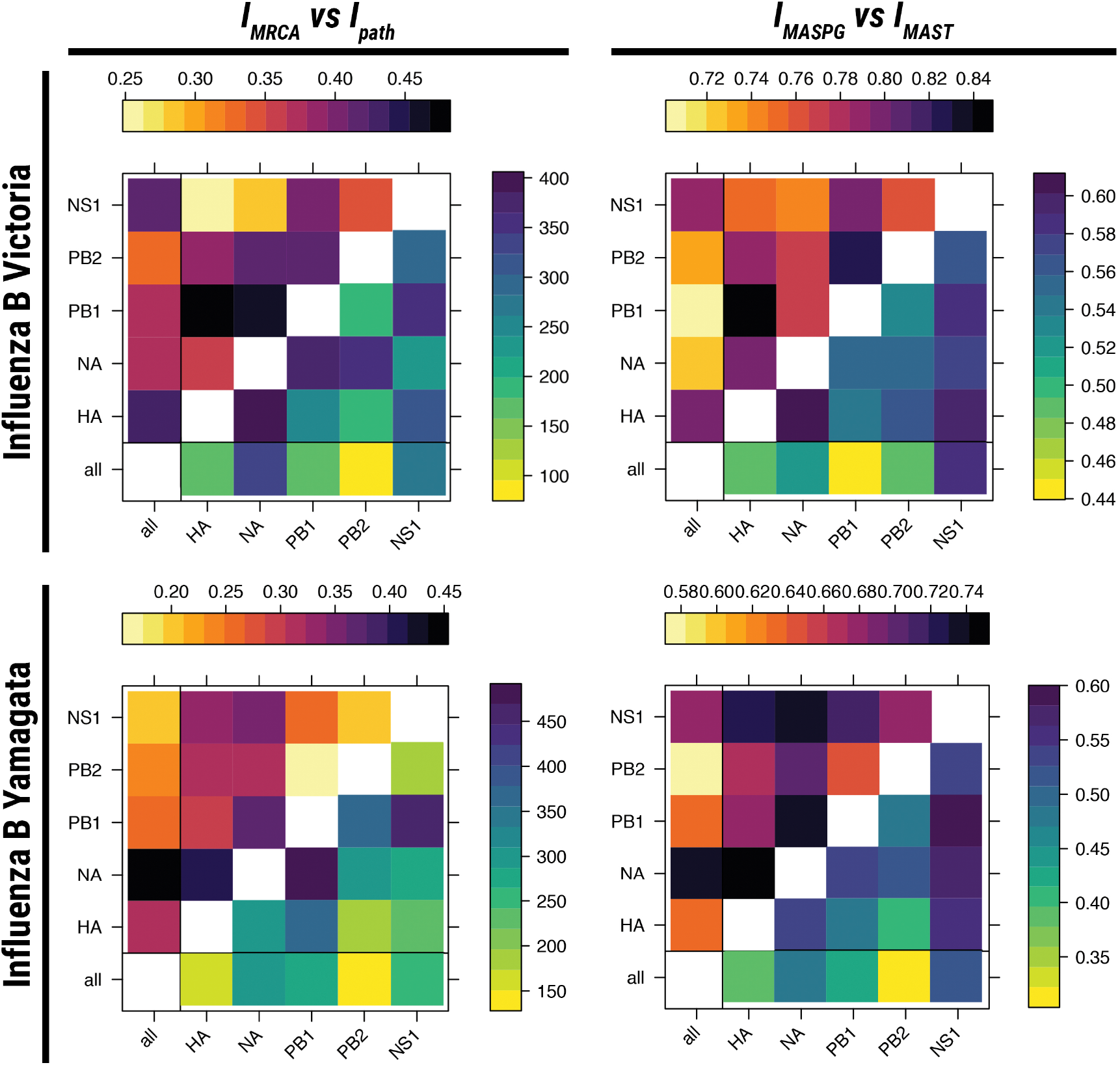
Comparisons of various incompatibility measures between phylogeographies reconstructed from segments of the influenza B genome for both the Victoria and Yamagata lineages. On the the left, *I_MRCA_* (upper triangle) is compared to *I_path_* (lower triangle) for the Victoria (upper plot) and Yamagata (lower plot) strains. On the right, *I_MASPG_* (upper triangle) is compared to *I_MAST_* (lower triangle) for the Victoria (upper plot) and Yamagata (lower plot) strains.

Despite free reassortment between all segments, the structure of incompatibilities follows some clear patterns. Phylogeographic incompatibilities between segments are stronger for the Victoria strain, while for Yamagata these are comparable to the ones of a recombining virus such as FMDV (see Figure 6). This is partly a result of the large impact of reassortment on the phylogenetic trees of the Victoria lineage.

There is a strong mismatch between the evolutionary histories of the two proteins that are the most important in the immune response against the virus (i.e. of HA and NA). The phylogenetic histories of HA and NA are highly divergent in the Victoria lineage, but the difference between their geographical histories is less extreme. On the other hand, the geographical spread of HA and NA in the Yamagata lineage followed very different routes, despite intermediate levels of reassortment. The geographical history of NA in this lineage is, in fact, markedly different from all other genes.

Clear patterns of incompatibility differ between the two lineages, as shown also in Figure 5. PB1 and PB2 phylogeographies are closely related for Yamagata but very different for Victoria. HA and NA phylogeographies are related to NS1 for Victoria, but the three genes present very different phylogeographies for Yamagata. Hence, the Victoria and Yamagata lineages differ not only in the tree topology, but also in the way different genes are spread geographically. For the Victoria lineage, different geographical histories in the basal lineages of the tree correspond to different movements among East Asian and Australasian regions, whilst in more recent years viruses are reconstructed to spread in different ways between Asia and Europe, with the trunk more concentrated in Asia for NA and HA and in Europe for the other segments. For the Yamagata lineage, whilst the clade 2 (i.e. B/Massachusetts/02/2012 clade) shows little evidence of incompatibility between geographical histories, the spatial evolution of different genomic segments along the trunk of clade 3 (i.e. B/Wisconsin/1/2010 clade) clearly follows different routes of epidemics across Europe, America and East Asia/Australia.

**Figure 5:**
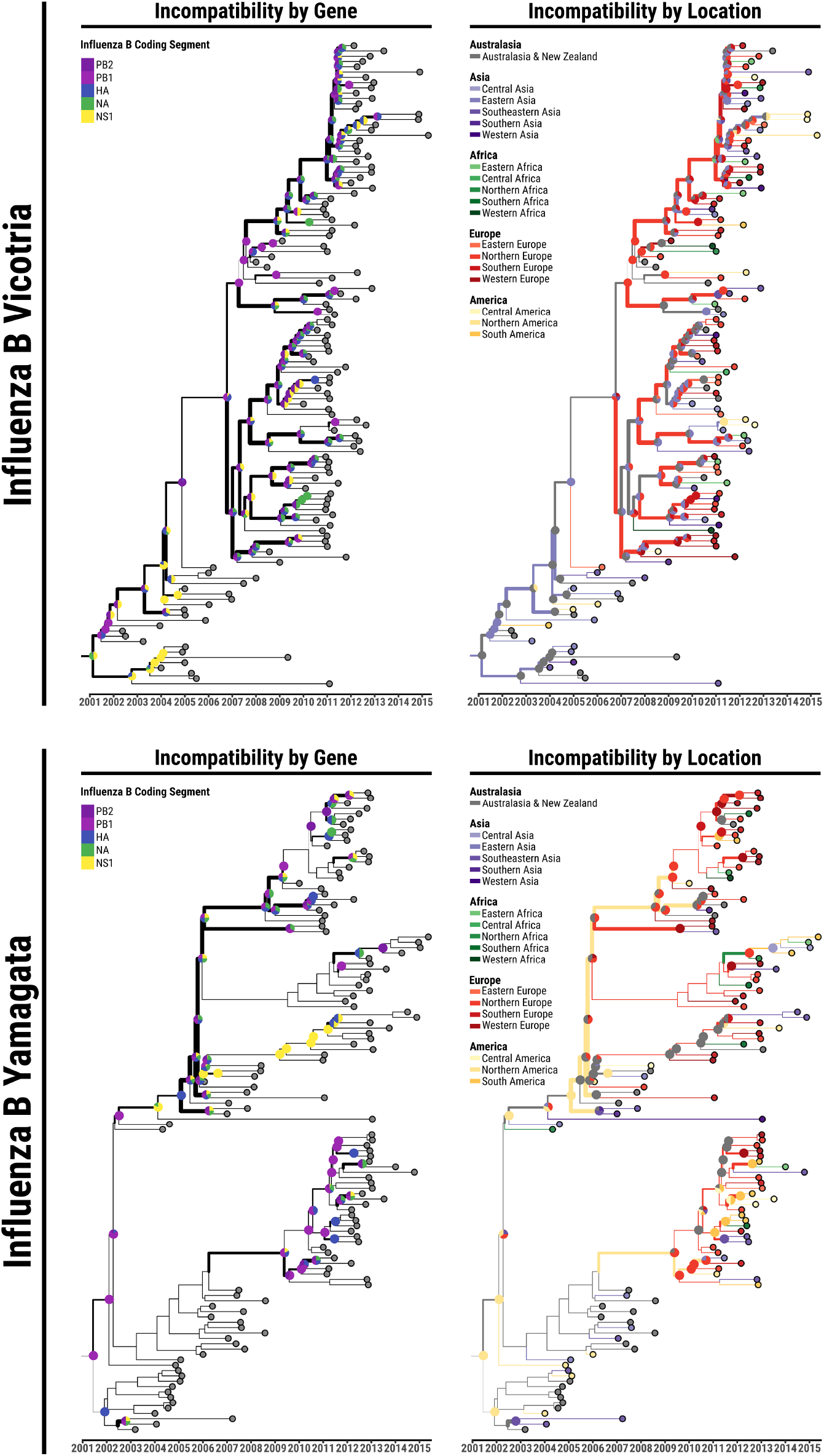
Illustration of the incompatibilities along the MCC tree for the Influenza B Victoria and Yamagata lineages epidemics. The width of each branch represents the total amount of phylogeographic MRCA incompatibilities between all trees built from individual genomic regions and the whole-genome tree. Pie charts epresent the distribution of MRCA incompatibilities per genomic region (left) and per location (right) on the corresponding nodes. Branch colors on the right represent the phylogeography reconstructed from the joint sequences alignment of all the genes investigated (~8.6 kb).

**Figure 6:**
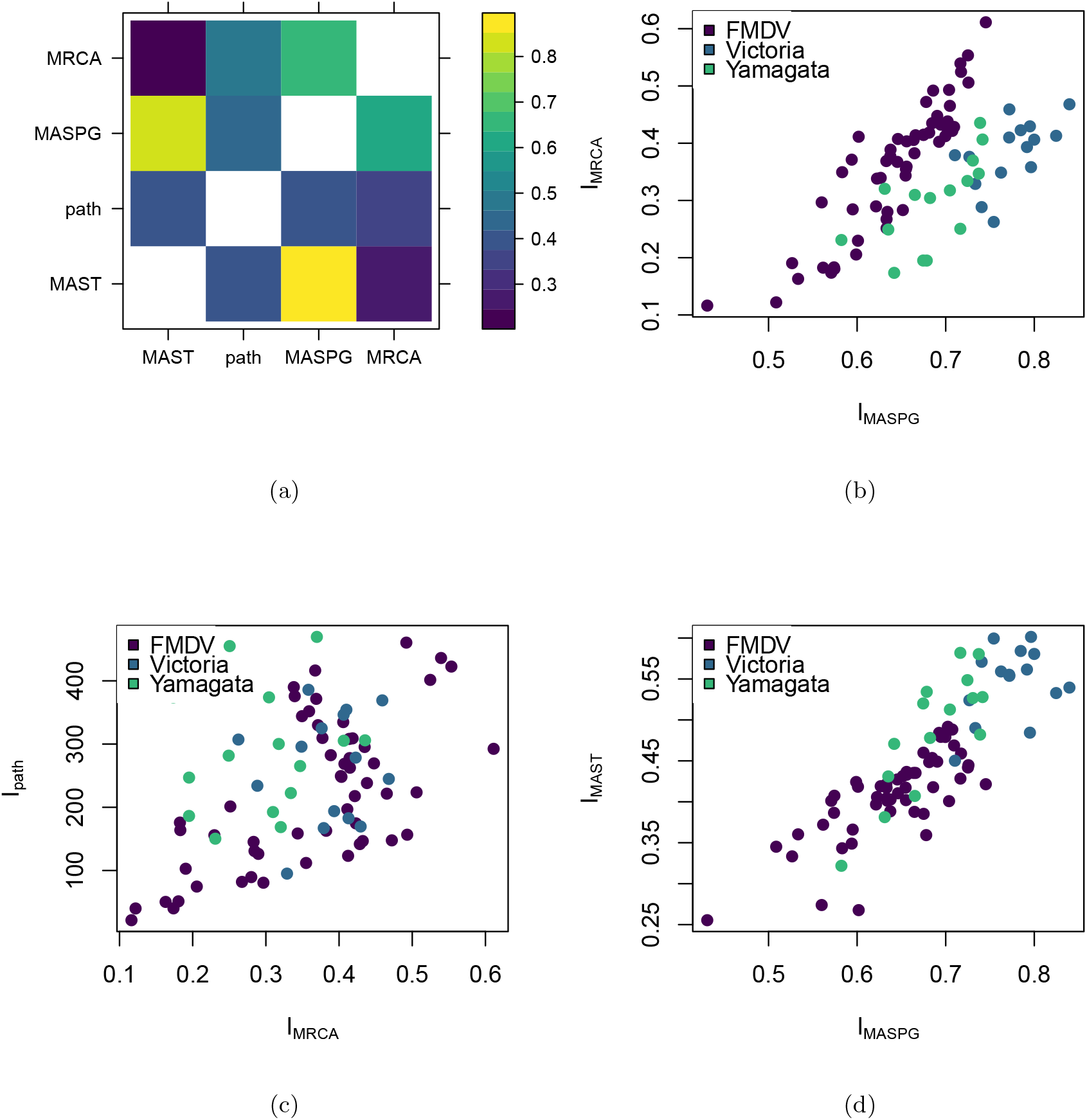
Illustrations of the correlations between the measures investigated in this paper. (a) Correlations between incompatibility measures. The upper triangle shows the Pearson correlation, and the lower triangle shows the Spearman correlation. (b-d) Scatter plots illustrating the correlations between incompatibility measures for the three viruses whose phylogeographies were investigated in this paper.

Also for the Influenza B case, the phylogeographic reconstruction from all genes together is not a reliable description of the evolutionary histories of individual genes. For both lineages, the overall phylogeny and phylogeography resembles mostly the one of PB2 and it is strongly incompatible with the geographic history of the NS1, hemagglutinin (in the Victoria lineage) and of the neuraminidase (in the Yamagata one).

## 4 Discussion

We have here presented two measures of phylogeographic concordance or incompatibility, *I*_*MRCA*_ and *I*_*MASPG*_. The first is derived from comparison of the geographical status of all the MRCAs of pairs of leaves. The second is derived from maximum agreement sub-phylogeographies. Both of these measures can be used to define metrics on the space of phylogeographies.

These incompatibility measures can be applied to any phylogeography, and more generally to any phylogeny where each tip has been assigned a geographical discrete location or any other continuous/discrete “trait” independent of the sequence. A typical example of such “traits” could be the host species for pathogens, or environmental variables for species living in a varied range of environments. The natural assignment method is parsimony as pioneered by Maddison, or Bayesian approaches such as BEAST [Suchard et al., 2018], but our measures are independent of the reconstruction method used and on the specific trait. Our approach can also compare multiple trees inferred from different origins, e.g. phylogenetic vs phylolinguistic trees, provided that the tips have the same geographical labels (representing e.g. different populations).

The two measures we propose are correlated but by no means identical. In fact, they are sensitive to different aspects of phylogeographic incompatibilities. We present these two particular measures in detail due to their naturalness and ease of computational implementation. There are however alternative possibilities, including extensions of tree distances other than the path and MAST distances. One promising such generalisation would be to include geographical information in Subtree-Prune-and-Regraft edit distances [De Oliveira Martins et al., 2014, Allen and Steel, 2001]. However, such a generalisation requires many arbitrary decisions in how internal node states are treated, and its values are challenging to compute. In practice, *I*_*MRCA*_ is both simple and sensitive enough to be the primary choice for phylogeographic incompatibilities.

There are obvious extensions to the present methods, such as allowing incomplete geographical metadata for the samples, constraints on the geographical assignments of internal nodes, or weighting geographical incompatibilities by their location on the phylogeny.

One important limitation of our approach is that we consider distinct phylogeographies, instead of a phylogeographic network or a geographically labelled ARG. This simplification is rooted in the current approaches to phylogeographic inference. In fact, both phylogenetic trees and phylogeographies are usually inferred from sequence data and geographical metadata; most often, inference is performed separately for each locus, resulting in posterior distributions with unknown correlations between loci. To account for uncertainties in the inference, we defined a measure of incompatibility over both posterior distributions of phylogeographies which is conservative with respect to potential correlations.

Eventually, a full stochastic model of sequence evolution, birth-death, transmission, recombination, and geography should be developed for joint inference at multiple loci. This is both conceptually and computationally hard, but in no way impossible. As a example, the model of [Dialdestoro et al., 2016] has these components except transmission and geography. Including transmission and geography is technically doable but the details of the implementation are challenging and it is not clear how it would scale to large data sets. Even analysing only intra-patient sequences, the model by [Dialdestoro et al., 2016] pushed the computational limits. Recent developments enable the inference of ARGs as collection of related trees for large genomes and large numbers of sequences [Kelleher et al., 2018], but it is unclear how the model can be extended to accommodate uncertainties and trait reconstruction. However, even a complete statistical model will not make the present work redundant, as any approach to statistical phylogeography could be faced with the need to define some kind of incompatibility between phylogeographies. In its simplicity, the approach outlined here is the first one to enable a systematic comparison and to provide a definition of distance between reconstructed phylogeographic histories.

## 5 Materials and Methods

### 5.1 Coalescent simulations

To investigate the dependence of *I*_*MRCA*_ on evolutionary parameters, we implemented a simulator of genealogies with migration and recombination. This simulator generates ancestral recombination graphs based on structured coalescent simulations in MASTER 6.1.1 [Vaughan and Drummond, 2013], implemented in BEAST2 2.6.1 [Bouckaert et al., 2019]. The simulation uses an island model of population structure with three populations of 20 individuals each and different patterns of migration between all populations (as illustrated in Figure 2). The simulations output geographically labelled ancestral recombination graphs (ARGs) for pairs of loci. We converted the ARGs into pairs of phylogeographies, one for each locus, and calculated their pairwise *I*_*MRCA*_ for each set of parameters. The mean *I*_*MRCA*_ for each parameter combination is shown in Figure 2.

### 5.2 Phylogeographic reconstruction

In order to test our method to real evolutionary scenarios, we compiled two different data sets of genome sequences from influenza B and foot-and-mouth disease (FMD) viral infections, which affect both human and livestock species. Reassortment events and recombination patterns have been described for both the viral infections along with their dynamics of geographical transitions [Carrillo et al., 2005, Jackson et al., 2007, Dudas et al., 2014, Bedford et al., 2015]. For influenza B virus we selected 122 and 120 unique genome sequences of, respectively, the Victoria (Vic) and Yamagata (Yam) lineages [Bedford et al., 2015, Langat et al., 2017], characterising five out of the eight gene segments and encoding: the polymerase basic subunits 1 and 2 (PB1 and PB2), the hemagglutinin (HA) and neuraminidase (NA) glycoproteins, and the non-structural protein 1 (NS1) [Bouvier and Palese, 2008]. For FMD we selected 74 whole-genome sequences (WGSs) characterising the O/ME-SA/Ind-2001 lineage [Bachanek-Bankowska et al., 2018], extracting separate alignments for each of the four structural (VP1 to VP4) and six non-structural (2A to 2C, and 3A to 3D) proteins, along with the leader polypeptide (Lpro) and the 5’ untranslated region (5’UTR) alignments [Jackson et al., 2003].

Distinct time-resolved phylogeographic trees were inferred for each of the influenza B genes and FMD virus (FMDV) proteins using BEAST 1.10.4 [Suchard et al., 2018]. Virus evolution was modelled by defining: the substitution process by the HKY [Hasegawa et al., 1985] and GTR+Γ4 [Tavaré, 1986] models respectively for influenza B and FMD; a log-normal relaxed molecular clock across branches [Drummond et al., 2006]; a flexible Bayesian Skyride coalescent prior on trees [Minin et al., 2008]. Phylogeographic patterns of lineage movement across geographic locations (defining discrete traits as countries for FMD, and as United Nations geoscheme regions for influenza B) were reconstructed using a discrete-state continuous-time Markov chain (CTMC) process, and assuming a non-reversible transition model with a Bayesian stochastic search variable selection (BSSVS) procedure [Lemey et al., 2009]. All priors were left at their default values. The Markov chain Monte Carlo (MCMC) chains were run for 200 million iterations, sampling trees every 20000 states. Inference were based on the resulting 9000 trees obtained after discarding the initial 10% of the chain as burn-in. Phylogeographic incompatibilities were assessed in the posterior phylogeographic space of a subset of 100 trees uniformly sampled from the posterior set of reconstructed phylogenies, using an approximation of the Wasserstein metric as described above.

## 6 Acknowledgements

We thank Nick Knowles and colleagues at the WRLFMD for useful discussions. The Pirbright Institute receives grant-aided support from the Biotechnology and Biological Sciences Research Council (BBSRC) of the United Kingdom (projects BBS/E/I/00007035, BBS/E/I/00007036 and BBS/E/I/00007039). This work was supported by funding from the BBSRC, grant number BBS/M011224/1. ADN acknowledges support from the Department for Environment, Food and Rural Affairs (Defra), United Kingdom, through funding from the SE2943 and SE2944 research grants.

## A Proofs that incompatibility measures are distances on 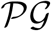

*I*_*MASPG*_ is a metric on 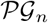. It is non-negative, and satisfies identity of indiscernibles and symmetry. The proof of the triangle inequality for *I*_*MASPG*_ is as follows.

Let us define *S*_*i*,*j*_ = *MASPG*(*T*_*i*_, *T*_*j*_). We require that, for three phylogeographies *T*_1_, *T*_2_, and *T*_3_,

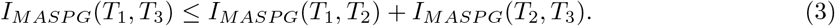

This is equivalent to requiring that

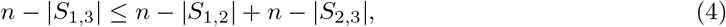

i.e.

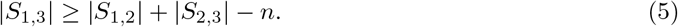

Let us note that any agreement sub-phylogeography between *S*_1,2_ and *S*_2,3_ must be a sub-phylogeography of all three trees, and so is a sub-phylogeography of *S*_1,3_. Let *U* be the largest geographically agreeing sub-phylogeography for all pairs of trees in {*T*_1_, *T*_2_, *T*_3_}. Now for each pair of trees the MASPG is either identical to *U* or is some larger agreement sub-phylogeography, so

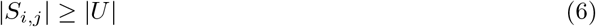

for all *i, j* ∈ {1, 2, 3}. Now, if *n* is the number of leaves, then *S*_1,2_ and *S*_2,3_ cannot share fewer than |*S*_1,2_| + |*S*_2,3_| − *n* leaves, since the union of the sets of leaves on the two sub-phylogeographies cannot be larger than *n*. Therefore

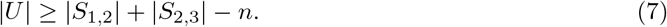

From (6) we get |*S*_1,3_| ≥ |*U*|, and substituting this into (7) yields (5), thus proving that the triangle inequality holds and that *I*_*MASPG*_ is a metric.

## B MASPG algorithm

A phylogeography, *P*_*a*_, is defined by a tree *T*_*a*_ and a mapping *f*_*G*_ from nodes of *T*_*a*_ to some geographic space 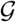. The algorithm for the MASPG of two phylogeographies is based on a modification of the Goddard algorithm for the MAST [Goddard et al., 1994]. The modification is based on this lemma, following [Rasmussen, 2016].

### Lemma 1.

*Let P*_*a*_ *be a phylogeography with tree T*_*a*_ *rooted at node a with the children b and c, and let P*_*w*_ *have a tree T*_*w*_ *rooted at w with children x and y*. *Let f*_*G*_(*a*) = *A*, *and f*_*G*_(*w*) = *W*. *We now define* #(*P*_*a*_, *P*_*w*_) *as the size of the maximum agreement sub-phylogeography (MASPG) of P*_*a*_ *and P*_*w*_. *If A* = *W*, #(*P*_*a*_, *P*_*w*_) *is given by the maximum of the following six numbers:* #(*P*_*b*_, *P*_*x*_)+#(*P*_*c*_, *P*_*y*_), #(*P*_*b*_, *P*_*y*_) + #(*P*_*c*_, *P*_*x*_), #(*P*_*a*_, *P*_*x*_), #(*P*_*a*_, *P*_*y*_), #(*P*_*b*_, *P*_*w*_), #(*P*_*c*_, *P*_*w*_). *If A* ≠ *W*, #(*P*_*a*_, *P*_*w*_) *is given by the maximum of* #(*P*_*a*_, *P*_*x*_), #(*P*_*a*_, *P*_*y*_), #(*P*_*b*_, *P*_*w*_), *and* #(*P*_*c*_, *P*_*w*_); *i.e. the last four numbers in the list above*.

*Proof.* This is a proof by case over the possible sub-phylogeographies contributing to the agreement sub-phylogeography *A*_*aw*_ for *P*_*a*_ and *P*_*w*_.

<u>Case 1:</u> Only *P*_*b*_ of *P*_*a*_ contributes to *A*_*aw*_. If this is the case, then the agreement sub-phylogeography must be an agreement sub-phylogeography of *P*_*b*_ and *P*_*w*_, and its size is equal to #(*P*_*b*_, *P*_*w*_).
<u>Case 2:</u> Only *P*_*c*_ of *P*_*a*_ contributes to *A*_*aw*_. If this is the case, then the agreement sub-phylogeography must be an agreement sub-phylogeography of *P*_*c*_ and *P*_*w*_, and its size is equal to #(*P*_*c*_, *P*_*w*_).
<u>Case 3:</u> Only *P*_*x*_ of *P*_*w*_ contributes to *A*_*aw*_. If this is the case, then the agreement sub-phylogeography must be an agreement sub-phylogeography of *P*_*x*_ and *P*_*a*_, and its size is equal to #(*P*_*a*_, *P*_*x*_).
<u>Case 4:</u> Only *P*_*y*_ of *P*_*w*_ contributes to *A*_*aw*_. If this is the case, then the agreement sub-phylogeography must be an agreement sub-phylogeography of *P*_*y*_ and *P*_*a*_, and its size is equal to #(*P*_*a*_, *P*_*y*_).
<u>Case 5:</u> All children *b*, *c*, *x*, and *y* contribute to the MASPG. It follows that *a* and *w* are part of the MASPG, and so *A* = *W*. Consider the ASPG *A*_*r*_, rooted in *r* with children *s* and *t*. We claim that a child of *r* cannot contain a leaf *b′* from *P*_*b*_ and a leaf *c′* from *P*_*c*_ at the same time. If this is the case, we only need to find the MASPG size of child pair permutations across *P*_*a*_ and *P*_*w*_ : *max* {#(*P*_*b*_, *P*_*x*_) + #(*P*_*c*_, *P*_*y*_), #(*P*_*b*_, *P*_*y*_) + #(*P*_*c*_, *P*_*x*_)}. The proof of this claim by contradiction is presented below.

**Figure B1.**
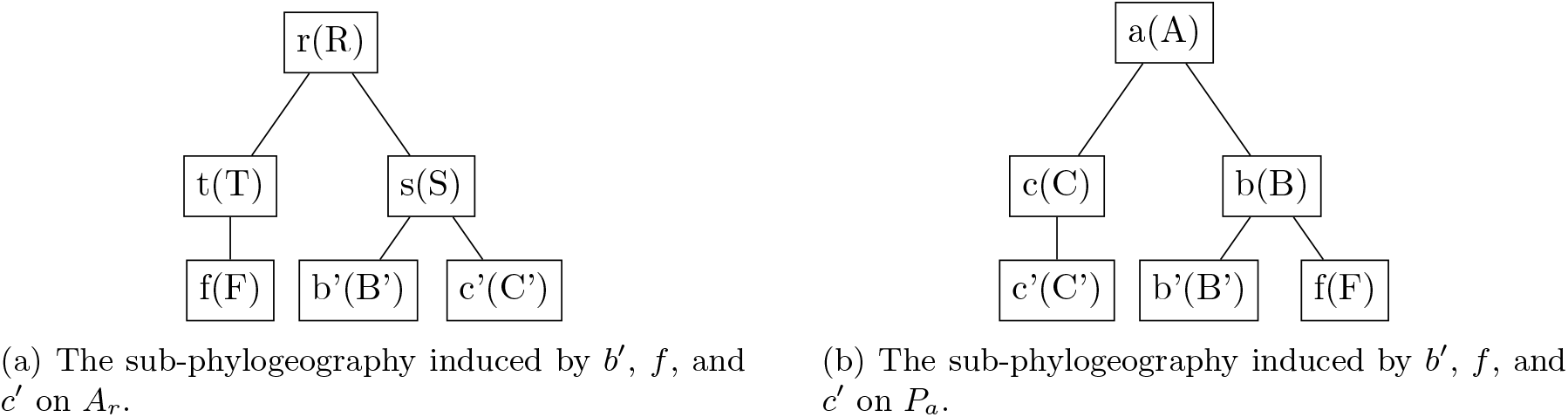
An illustration of the central claim in Case 5 of the proof by case of Lemma 1. The figures show that a child of *r* cannot contain a leaf *b′* from *P*_*b*_ and a leaf *c′* from *P*_*c*_ at the same time, since the sub-phylogeographies induced by these leaves and a leaf from a child of *r* on *A*_*r*_ and *P*_*a*_ must differ topologically.

Consider the ASPG *A*_*r*_, rooted in *r* with children *s* and *t*. Suppose that *A*_*s*_ contains a leaf *b′* from *P*_*b*_ and a leaf *c′* from *P*_*c*_. Let *f* be any leaf from *A*_*t*_. Assume that *f* is from *P*_*b*_ (the proof for the other case is equivalent). Then the sub-phylogeography induced by *b′*, *f*, and *c′* must differ for *A*_*r*_ and *P*_*a*_, as illustrated in Figure 1.

The algorithm to calculate *I*_*MASPG*_ can then be constructed by iterating the process of finding the maximum of the sizes of various agreement sub-phylogeographies, working from the leaves upwards.

**Figure B2:**
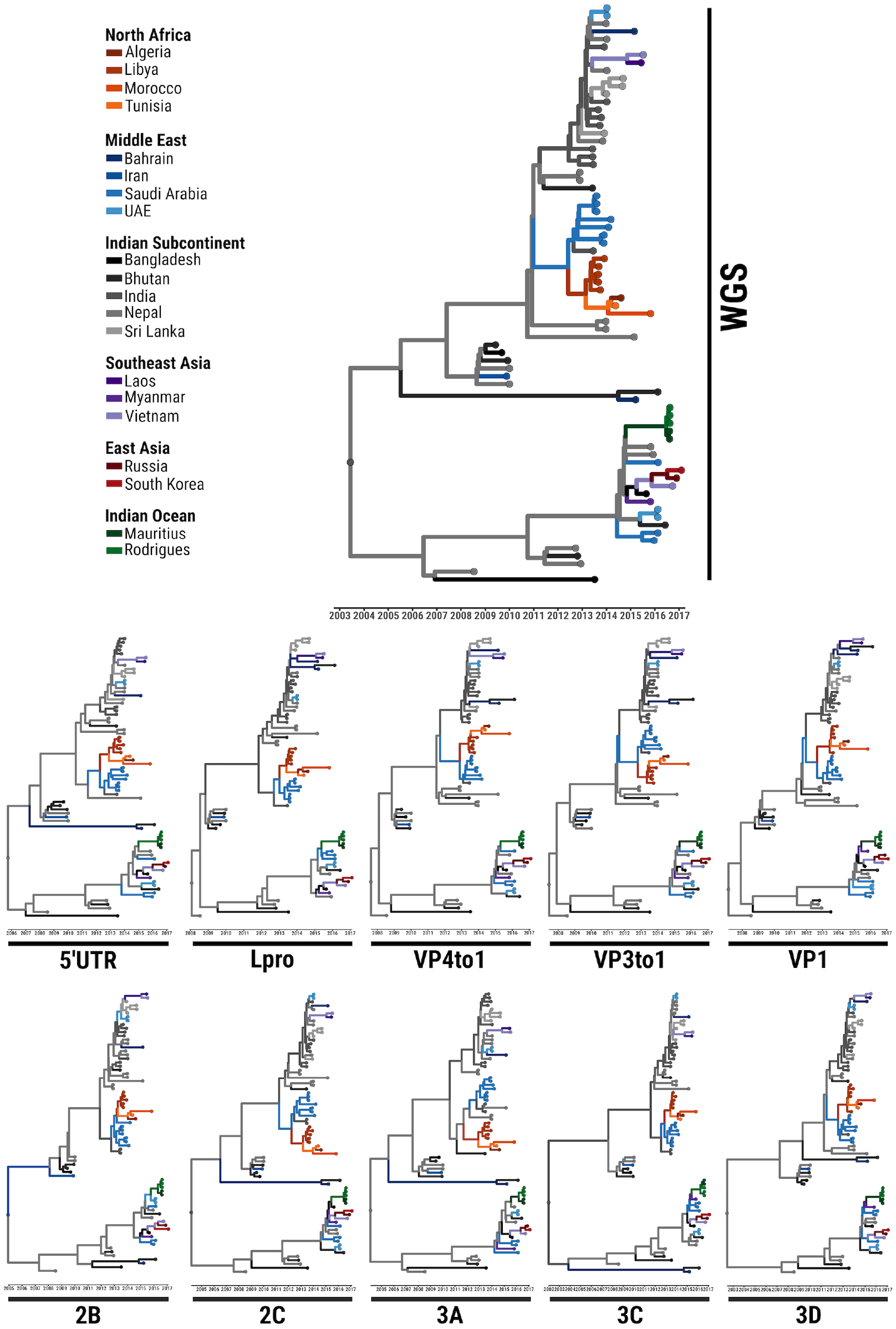
MCC phylogeographies of the O/ME-SA/Ind-2001 epidemic reconstructed for the whole-genome sequence and each of the FMDV protein.

**Figure B3:**
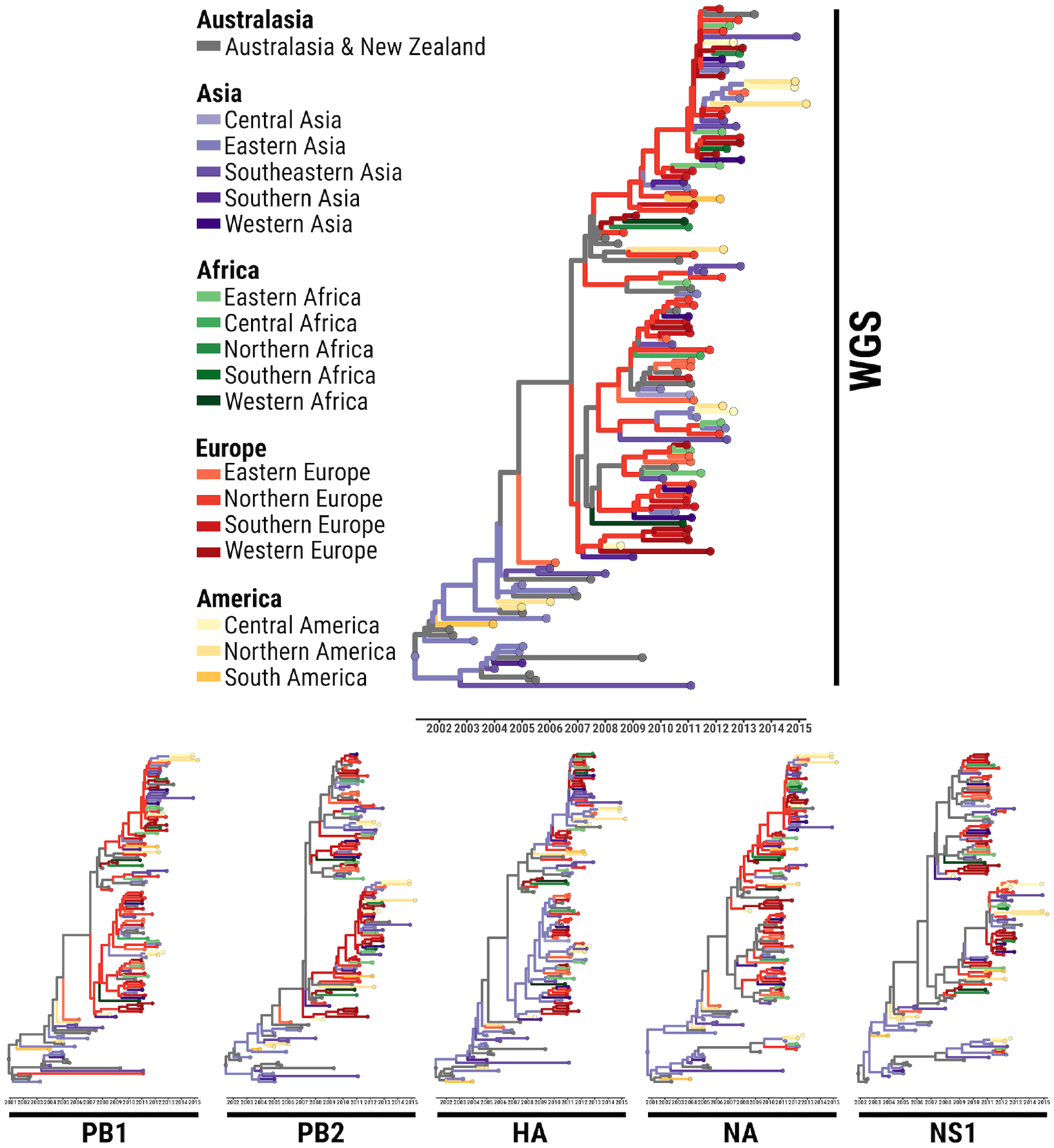
MCC phylogeographies of the Influenza B Victoria epidemics reconstructed for the whole-genome sequence and each of the segment.

**Figure B4:**
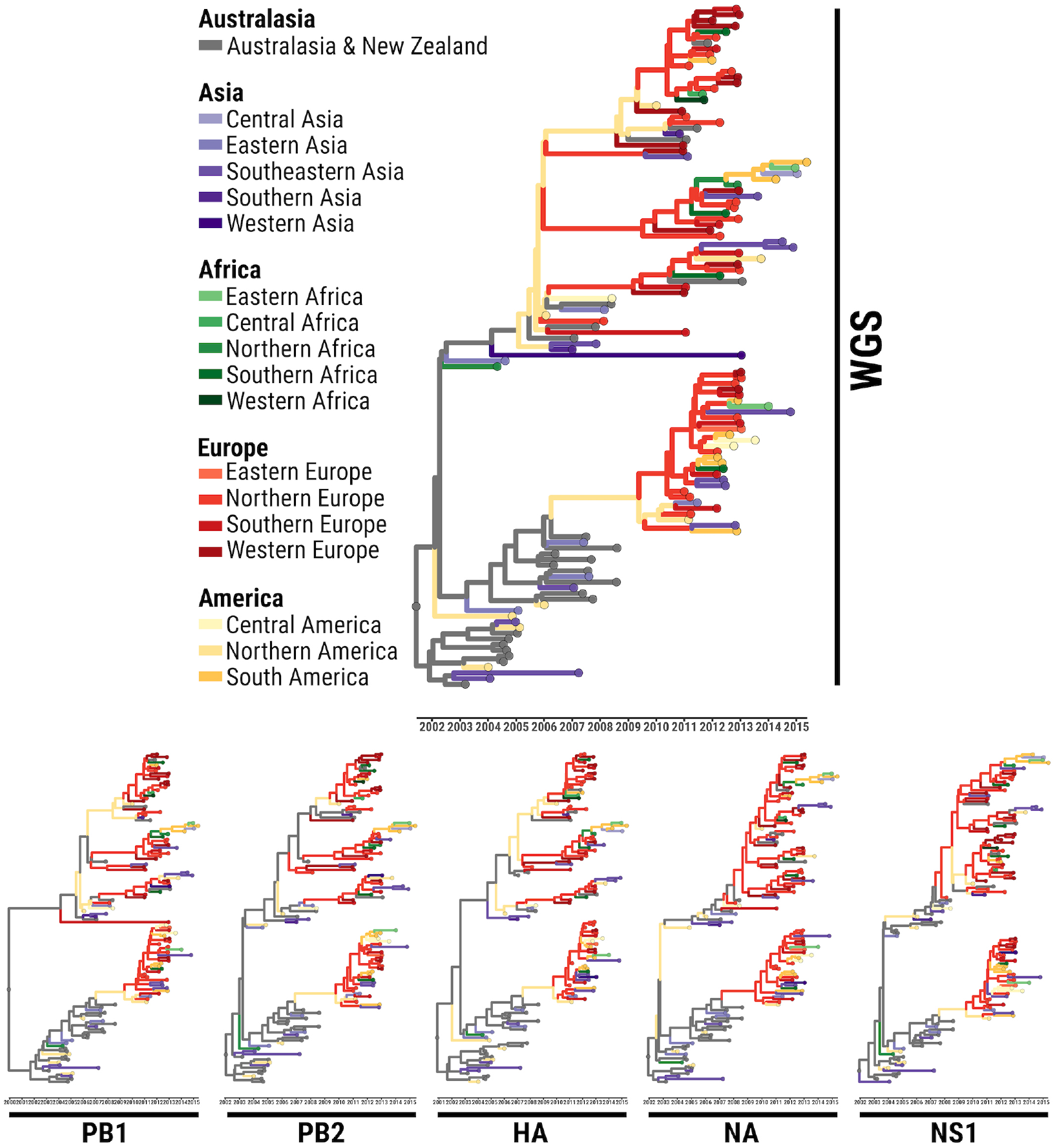
MCC phylogeographies of the Influenza B Yamagata epidemics reconstructed for the whole-genome sequence and each of the segment.

**Figure B5:**
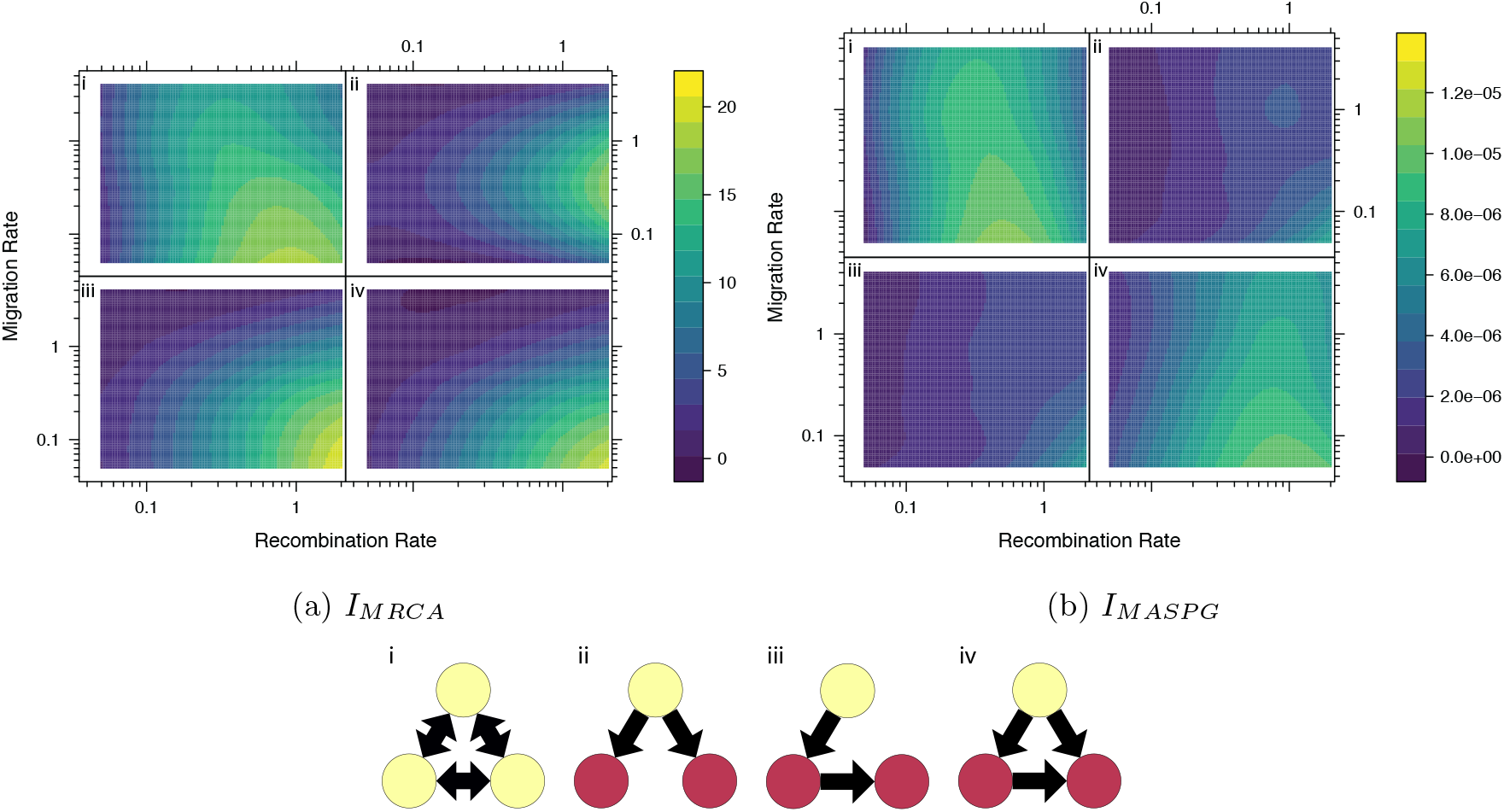
Effect of evolutionary parameters on the variance of phylogeographic incompatibilities, using 100 simulations at each calculated pair of parameter values. The plots show smoothed surfaces of the variance each phylogeographic incompatibility measure, labelled with the corresponding network structure. The network structures are illustrated with locations in which the population can originate coloured yellow.

**Figure B6:**
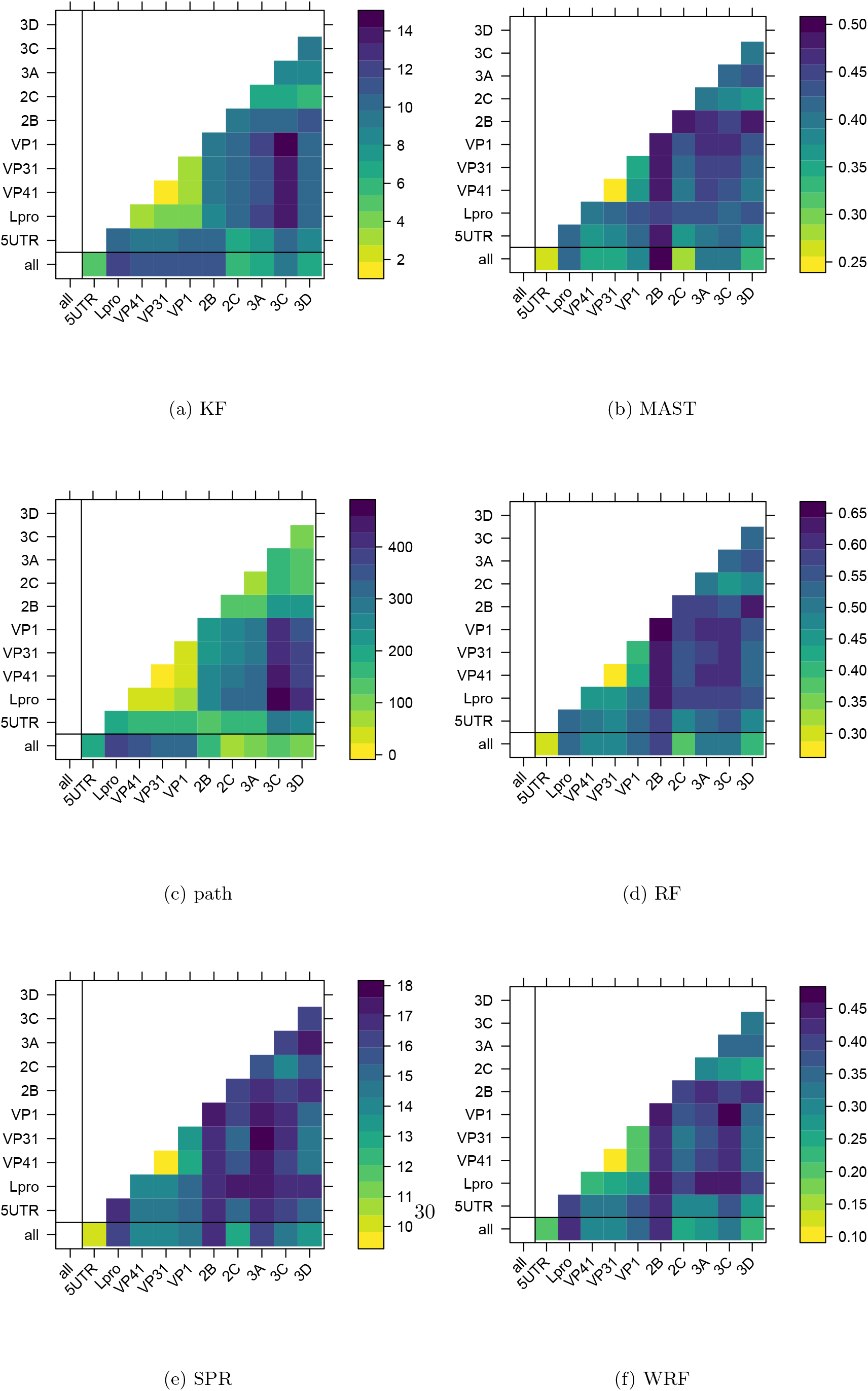
Phylogenetic incompatibilities across genomic regions in FMDV

**Figure B7:**
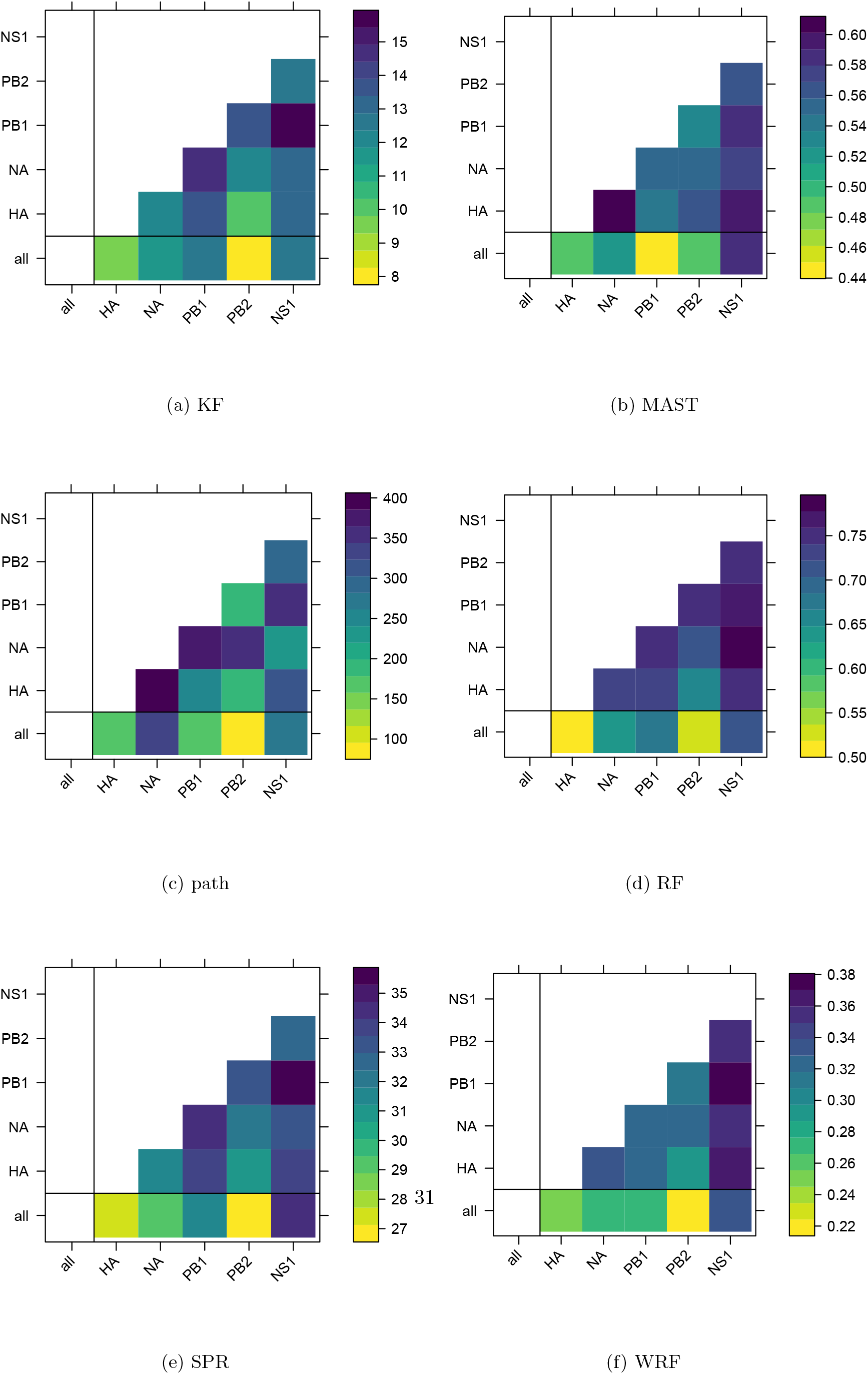
Phylogenetic incompatibilities across genomic regions in Influenza B Victoria

**Figure B8:**
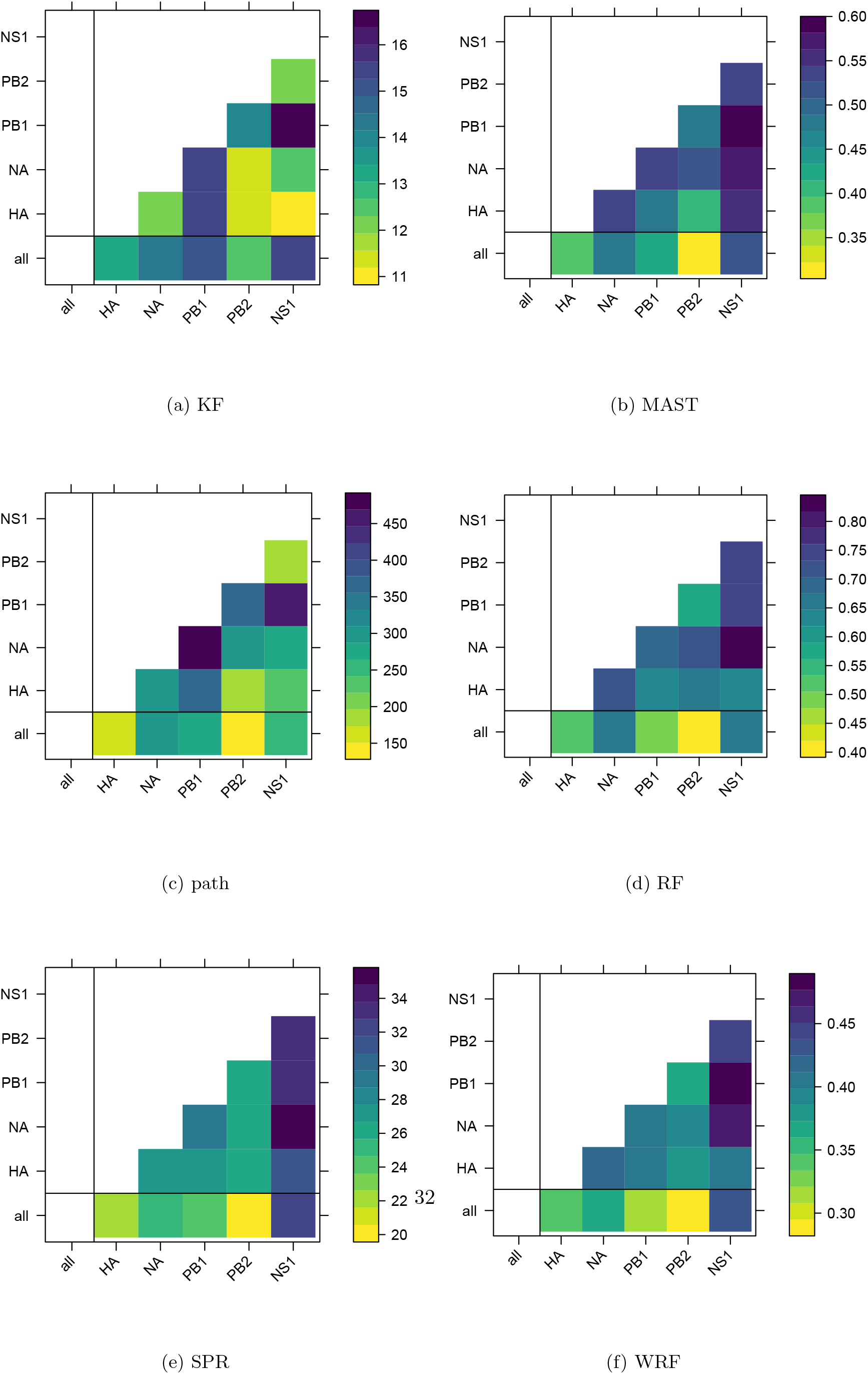
Phylogenetic incompatibilities across genomic regions in Influenza B Yamagata

